# Incorporation of native HCV E1E2 into a nanoparticle vaccine platform

**DOI:** 10.64898/2026.08.05.743069

**Authors:** Liudmila Kulakova, Sarah Jeong, DongXiu Zhang, Xiaoran Shang, Kinlin L. Chao, Alexander Marin, Matthew C. Metcalf, Tagide deCarvalho, Edwin Pozharski, Yuxing Li, Brian G. Pierce, Alexander K. Andrianov, Eric A. Toth, Thomas R. Fuerst

## Abstract

Development of an effective HCV vaccine requires the induction of both broadly neutralizing antibodies (bnAbs) and a robust cellular response. One issue that has arisen is that HCV subunit vaccines have limited immunogenicity, thus requiring multivalent formats in order to elicit a robust anti-HCV immune response. Toward that end, nanoparticle vaccines possess the ability to facilitate a controlled multivalent presentation and trafficking to lymph nodes, where they can interact with both arms of the immune system. Here, we used a soluble, secreted form of E1E2 (sE1E2) to assemble native E1E2 into a nanoparticle platform using a post-purification coupling assembly system. Nanoparticles were assembled by purifying sE1E2 containing a C-terminal SpyTag and an mi3-SpyCatcher fusion separately and covalently coupling the components via incubation. Free sE1E2-SpyTag was removed from nanoparticle preparations via gel filtration. The sE1E2-mi3 nanoparticles are fully competent to bind conformation-dependent bnAbs, indicating retention of a native assembly in the nanoparticle format. Electron microscopy analysis showed a clear incorporation of sE1E2 on the surface of the nanoparticle. Immunogenicity of sE1E2-mi3 nanoparticles was examined relative to sE1E2 alone and membrane-bound E1E2 (mbE1E2) following inoculation of groups of CD1 mice. Assessment of the immunogenicity of the sE1E2-mi3 nanoparticles showed that the nanoparticle assembly has a similar immunogenicity profile to that of mbE1E2 after only a prime and one boost, and overall superior to sE1E2. This proof-of-principle study sets the stage for further exploration of nanoparticles and other multivalent platforms for the development of E1E2-based vaccines.

**Importance:** Hepatitis C virus infects approximately 50 million people, and at present no effective HCV vaccine exists. Due to the high sequence variability of HCV and the resulting difficulty in developing a vaccine that elicits a broadly neutralizing response, multiple efforts are underway to enhance the immunogenicity of HCV vaccine candidates. In this study, we incorporated native soluble, secreted E1E2 (sE1E2) into a 60-mer nanoparticle via the SpyTag-SpyCatcher system and covalent isopeptide bond attachment using the purified components. These nanoparticles are antigenically intact and elicit a neutralizing antibody response at an earlier time point in the immunization regimen than the corresponding subunit vaccine. These studies show that a well-characterized sE1E2 platform compatible with multiple genotypes can be coupled to nanoparticles for use as a vaccine candidate.

## Introduction

Subunit vaccines comprise a portion of the infectious agent that is known to be immunologically important and are typically safer to use than live, attenuated organisms that have been used over the course of vaccination history. This enhanced safety stems in part from the ability to produce highly purified preparations of the subunit vaccine using recombinant expression platforms (1). However, the ease of production and high level of purity comes at the expense of immunogenicity. One factor is size, which affects antigen uptake and clearance/degradation, and it also determines the presentation mode of the vaccine antigen. Subunit vaccines are most often either monomeric or low-order oligomers (e.g. HIV env trimers (2)), whereas live and attenuated viruses present the vaccine antigen in a polyvalent manner in a confined volume on the virion itself. This effect has been observed for inactivated HCV produced in Huh7.5 hepatoma cells (3), where the neutralization titer achieved after immunization of mice with inactivated HCV was significantly enhanced relative to that for mice immunized with the E2 ectodomain alone.

Particulate antigens are generally known to be highly immunogenic (4–8). In a study by Aung et al. (9), the authors showed that subunit vaccines were trafficked primarily to the subcapsular sinus or extracellular regions of lymph nodes and subsequently degraded by metalloproteases, eliminating conformation-dependent epitopes on the associated antigens and hampering the immune response. Nanoparticle-sized antigens were localized instead to follicular dendritic cells (FDCs) where they remained intact and preserved such conformation-dependent epitopes and thus elicited a more robust immune response. In light of these observations, increasing the size of a subunit vaccine should be beneficial. A common strategy to achieve this end is via nanoparticle platforms, which are typically naturally occurring or engineered protein shells that allow for the multivalent display of subunit vaccines on the exterior (10–16). These assemblies can be formed in cis, where the subunit vaccine and nanoparticle protomer are expressed as a single open reading frame and assembly yields a 100% occupied nanoparticle, or in trans where the nanoparticle shell and subunit vaccine are produced separately and coupled post hoc, as in the case of the SpyCatcher-SpyTag system (17). Several platforms have been explored for a potential HCV vaccine candidate (18–20). The first studies by Yan et al. (20) and He et al. (18) used the E2 ectodomain and a modified E2 core ectodomain, respectively, as proof-of-principle antigens to be appended to nanoparticles. Given the importance of the E1E2 complex, and in particular the AR4/AR5 antigenic region in viral clearance (21, 22), a nanoparticle-presenting native E1E2 should be a high priority for HCV vaccine development. One nanoparticle study has been conducted with E1 and E2 (19), but this used a permuted E2-E1 version of the antigen which does not retain the native AR4/AR5 antigenic domain. In addition, we recently collaborated on a study which successfully produced an in cis E1E2 nanoparticle vaccine candidate (23). In the present study, we have employed the trans nanoparticle platform with well-characterized soluble secreted E1E2 heterodimers (24, 25).

HCV is a major cause of severe liver diseases and cancer, and the global burden is over 50 million chronically infected individuals with an annual increase of 1 million new infections (26). HCV infection progresses to chronic illness in nearly 75% of cases, which markedly increases the risk for development of cirrhosis or hepatocellular carcinoma. The World Health Organization recently announced a global hepatitis strategy, which aims to reduce new infections from all types of hepatitis viruses by 90% and associated deaths by 65% by 2030 (27). Direct-acting antivirals (DAAs), which cure existing HCV infections, (28, 29) are a major component of this strategy. However, DAA-treated individuals can become re-infected which decreases effectiveness in high-risk groups. Moreover, most HCV-infected individuals are asymptomatic until liver damage is extensive. Therefore, there is a clear need for an effective HCV vaccine (30) which would complement the use of DAAs. The high degree of genetic diversity within the seven major genotypes of HCV makes developing a vaccine for this pathogen challenging. However, spontaneous clearance rates and the presence of immune memory among individuals who clear their first HCV infection (31–34) suggest that development of an HCV vaccine is feasible (30, 35–37). Leveraging a flexible platform such as protein-based nanoparticles would likely improve prospects for HCV vaccine development.

In this study, we established a method for incorporating the well-characterized soluble secreted E1E2 antigen (sE1E2.SZ (25)) into a plug-and-display (17) mi3 nanoparticle system. Analysis by negative stain electron microscopy (EM) clearly shows incorporation of the sE1E2.SZ heterodimer on the outside of the nanoparticle shell and analytical ultracentrifugation shows no evidence of unoccupied nanoparticles using our protocol. The nanoparticles are antigenically intact, exhibiting only modest differences in affinity relative to the sE1E2.SZ heterodimer. We immunized mice with these nanoparticles and assessed the immune response relative to the sE1E2 subunit vaccine and membrane-bound E1E2 (mbE1E2) controls. An antibody response to immunization develops earlier in the immunization regimen (day 14) for mice immunized with the nanoparticles and mice immunized with mbE1E2, which is known to form large agglomerates containing multiple mbE1E2s (38), whereas the subunit vaccine groups do not show a detectable antibody response until day 28. Similarly, a neutralizing antibody response against the homologous pseudovirus is robust at day 28 for mice immunized with the nanoparticles and mice immunized with mbE1E2 but is comparably weaker for the subunit vaccine groups. Moreover, we observe cross-neutralization against a genotype 1b pseudovirus for the group immunized with sE1E2.SZ-containing nanoparticles and mbE1E2 at day 28. These results demonstrate that the sE1E2.SZ antigen can be incorporated into a plug-and-display nanoparticle platform, opening up the possibility of developing mosaic HCV vaccines in a similar fashion to those against SARS-CoV2 (39–41) and influenza (42–44).

## Results

### Nanoparticle strategy

For our proof-of-principle study, we chose to use the well-characterized soluble, secreted E1E2 (sE1E2) containing a synthetic leucine zipper dimeric coiled coil (SYNZIP1/SYNZIP2, or SZ) to be coupled to mi3-based nanoparticles (11) via SpyTag-SpyCatcher (45) covalent attachment (**Fig. 1A**). This allowed us to perform a side-by-side comparison between the identical subunit vaccine (sE1E2.SZ) and the nanoparticle (sE1E2.SZ-mi3). To accomplish this, we synthesized a version of sE1E2.SZ with an optimized SpyTag (45) appended to the C-terminus of the SYNZIP1 scaffold appended to the E2 ectodomain (sE1E2.SZ.SpyTag; for simplicity we refer to this as sE1E2.SZ throughout the manuscript). For the nanoparticle itself, based on previous data pertaining to sE2 ectodomain nanoparticles (18), we chose the mi3 nanoparticle with a corresponding optimized SpyCatcher appended to the N-terminus of the mi3 protomer (SpyCatcher-mi3). These components were expressed, purified, and coupled together to form the sE1E2.SZ-mi3 nanoparticle as described below.

**Figure 1.**
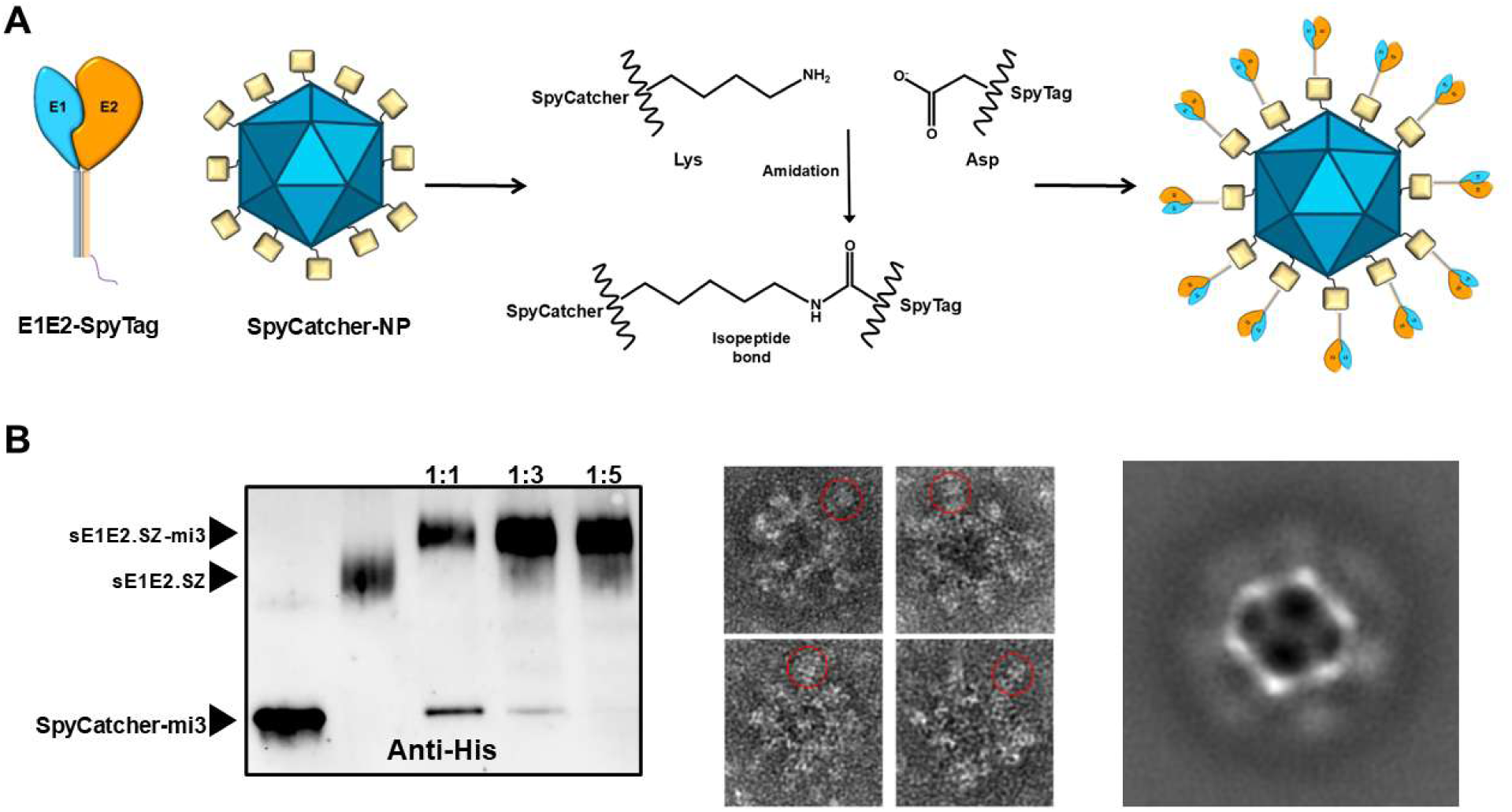
**A.** Schematic of sE1E2.SZ-mi3 nanoparticle assembly *in trans*. **B**. A Pilot scale coupling monitored by western blot and nsEM. (left) Western blot probed with an anti-His antibody showing a shift of SpyCatcher-mi3 to a higher molecular weight upon incubation of different ratios of sE1E2.SZ.SpyTag. (middle) Individual micrographs showing sE1E2.SZ decorating the SpyCatcher-mi3 nanoparticle shell as indicated by red circles. (right) Class average of 308 particles showing string density for the mi3 nanoparticle shell and diffuse density for the attached sE1E2.SZ.SpyTag.

### Coupling and purification of NPs

Purified sE1E2.SZ and SpyCatcher-mi3 were first incubated on a small scale at varying ratios of sE1E2.SZ to SpyCatcher-mi3 to assess coupling efficiency. This pilot scale coupling was analyzed by negative stain electron microscopy to confirm sE1E2 incorporation (described below). Based on this pilot experiment (**Fig. 1B**), a molar ratio of 5 sE1E2.SZ per Spycatcher-mi3 was chosen for larger scale coupling in preparation for bioanalytical analysis and animal immunization. For larger scale coupling, 6.8 mg of sE1E2.SZ and 2.66 mg of SpyCatcher-mi3 were incubated for 20 h at 4 °C. Following incubation, coupled sE1E2.SZ-mi3 nanoparticles were purified from uncoupled components by gel filtration using a HiPrep 16/60 Sephacryl S-500HR column. The final coupled material, along with the other antigens used in this study (sE1E2.SZ, mbE1E2, and mi3) were analyzed by SDS-PAGE and both reducing and non-reducing western blot (**Fig. 2**).

**Figure 2.**
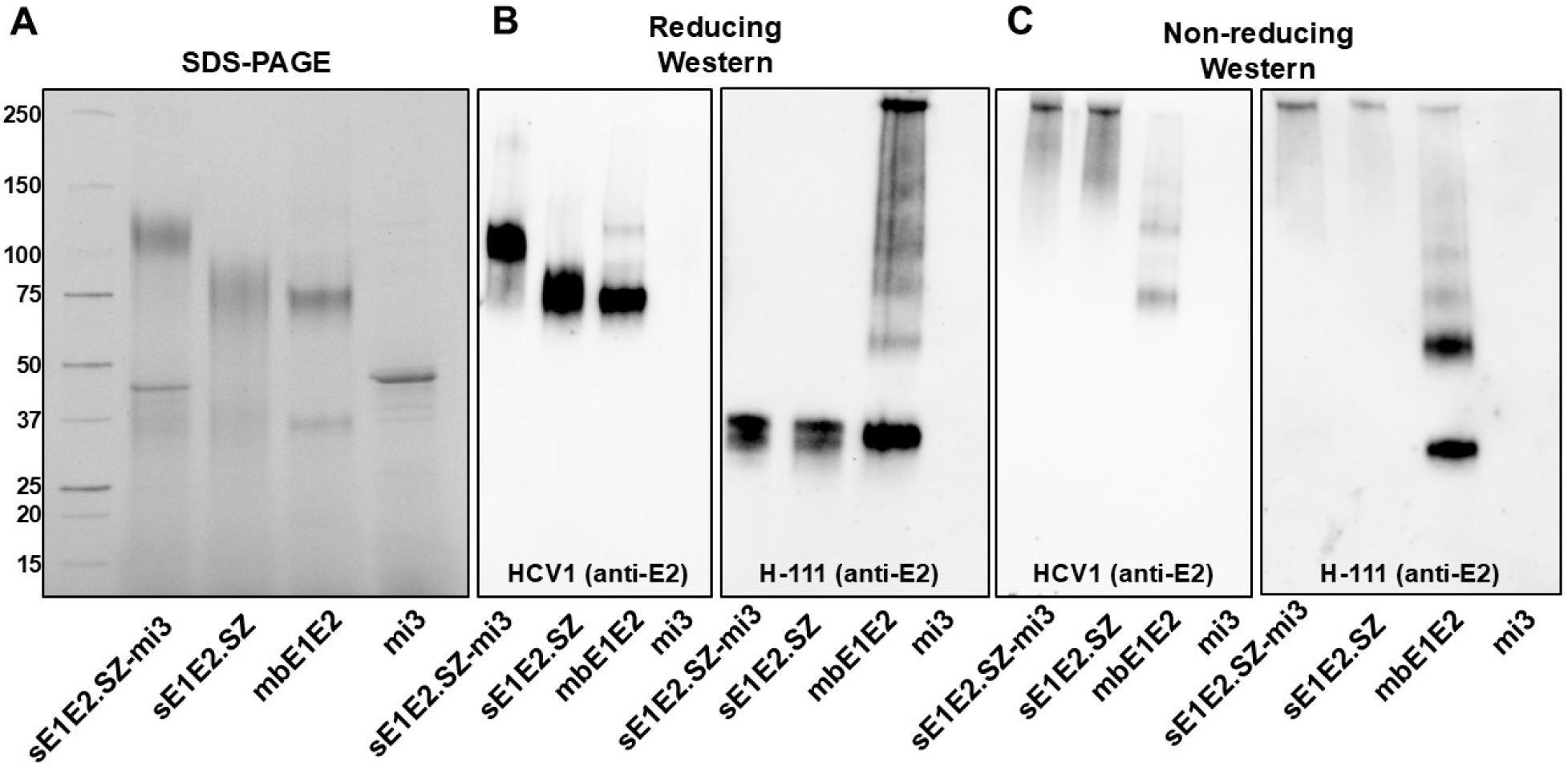
Purified antigen protein analysis. **A.** SDS-PAGE analysis of purified antigen sE1E2.SZ-mi3, sE1E2.SZ, mbE1E2, and mi3 under reducing conditions. **B.** Western blot detection using the anti-HCV E2 antibody HCV1 and the anti-HCV E1 antibody H-111 under reducing conditions. **C.** Western blot detection using the anti-HCV E2 antibody HCV1 and the anti-HCV E1 antibody H-111 under non-reducing conditions.

### Negative stain *EM*

Pilot scale sE1E2.SZ-mi3 nanoparticles were analyzed at the Keith R. Porter imaging facility at the University of Maryland Baltimore County. Analysis of individual particles from the pilot study showed clear incorporation of sE1E2 (red circles, **Fig. 1B**). A class average of 308 particles from this experiment also shows the core nanoparticle shell decorated with additional density of varying intensity (**Fig. 1B**). The sE1E2.SZ-mi3 nanoparticles used for bioanalytical characterization and animal immunization were further characterized by negative stain electron microscopy at the Maryland Center for Advanced Molecular Analysis (M-CAMA). Both control mi3 nanoparticles and sE1E2.SZ-mi3 nanoparticles were imaged on an FEI Talos Arctica 200 kV TEM equipped with a FEI Falcon3EC detector, and data were collected using SerialEM and processed using cryoSPARC. The sE1E2.SZ-mi3 nanoparticle class averages show clear decoration of varying intensity on the surface of the mi3 nanoparticle shell (**Fig. 3A**). By contrast, the control mi3 nanoparticle shows strong density for the nanoparticle shell both in 2D class averages and in the 3D reconstruction (**Fig. 3B**) with no extraneous density. Consistent with this, the 3D reconstruction of sE1E2-mi3 shows a well-defined mi3 nanoparticle shell with diffuse electron density protruding from the surface. This additional density, while not well-defined enough to accurately place a model, extends far enough from the surface to accommodate sE1E2, indicating that the larger scale coupling was successful in generating sE1E2.SZ coupled to SpyCatcher-mi3.

**Figure 3.**
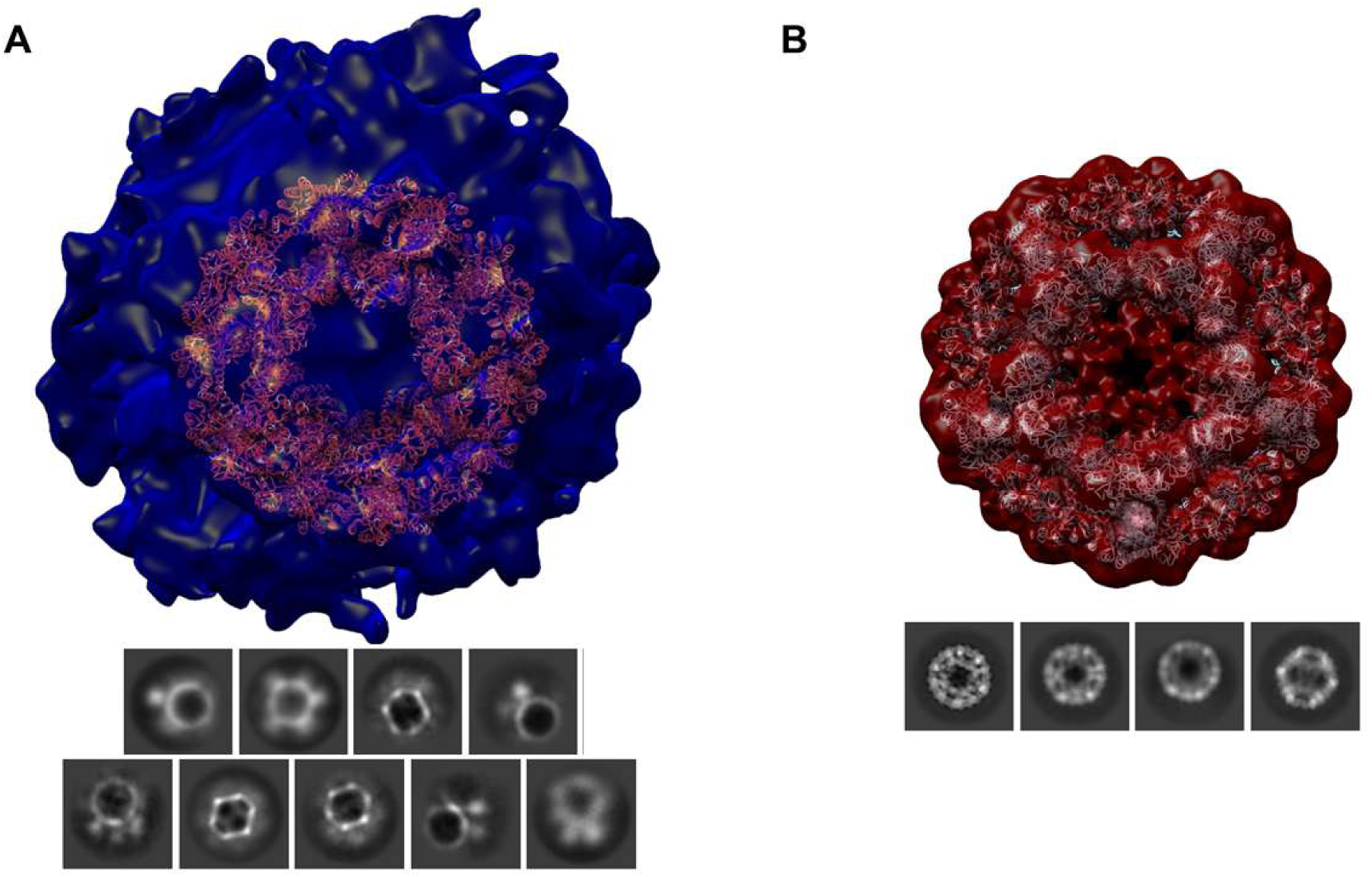
Analysis of sE1E2.SZ-mi3 and mi3 nanoparticles by nsEM. **A.** (top) 3D reconstruction of the sE1E2.SZ-mi3 nanoparticle calculated using CryoSPARC with no symmetry imposed. A docked model of mi3 (RCSB ID 5kp9) is shown in orange. Clear additional density on the surface indicative of coupled E1E2, but the density is not distinct. (bottom) 2D class averages from the sE1E2.SZ-mi3 nanoparticle sample. The nanoparticle backbone can be distinguished along with sometimes diffuse, sometimes distinct E1E2 density on the surface. B. (top) 3D reconstruction of the mi3 nanoparticle calculated using CryoSPARC with icosahedral symmetry imposed. Docked model of mi3 (RCSB ID 5kp9) shows excellent agreement with EM density. (bottom) 2D class averages from the mi3 nanoparticle sample. The nanoparticle backbone is readily distinguishable in the class averages.

### Analytical characterization of solution heterogeneity

We used analytical ultracentrifugation (AUC) to assess the size distribution of the sE1E2.SZ-mi3 nanoparticles used for animal immunization relative to control mi3 nanoparticles and sE1E2.SZ samples. AUC can separate a mixture of protein populations more precisely than SEC (82), and thus our aim was to ascertain whether any naked nanoparticles remained in the sE1E2.SZ-mi3 preparations. These particles sediment at values consistent with very large assemblies, with a discrete peak at 61 S and a minor peak at 90 S (**Fig. 4A**). These two peaks could represent different levels of incorporation of sE1E2 into the nanoparticle or a population that contains some fraction of dimeric sE1E2.SZ incorporated into the nanoparticle. By contrast, the mi3 alone control contains a main species of sedimentation coefficient 32 S with a very minor species at 48 S (**Fig. 4B**). Neither of these peaks overlaps with those of the sE1E2.SZ-mi3 nanoparticles, indicating that there are no detectable unoccupied particles in the sE1E2.SZ-mi3 preparations. Similarly, the control sE1E2.SZ samples show two species that migrate at significantly lower S values than either of the nanoparticle samples (**Fig. S1**).

**Figure 4.**
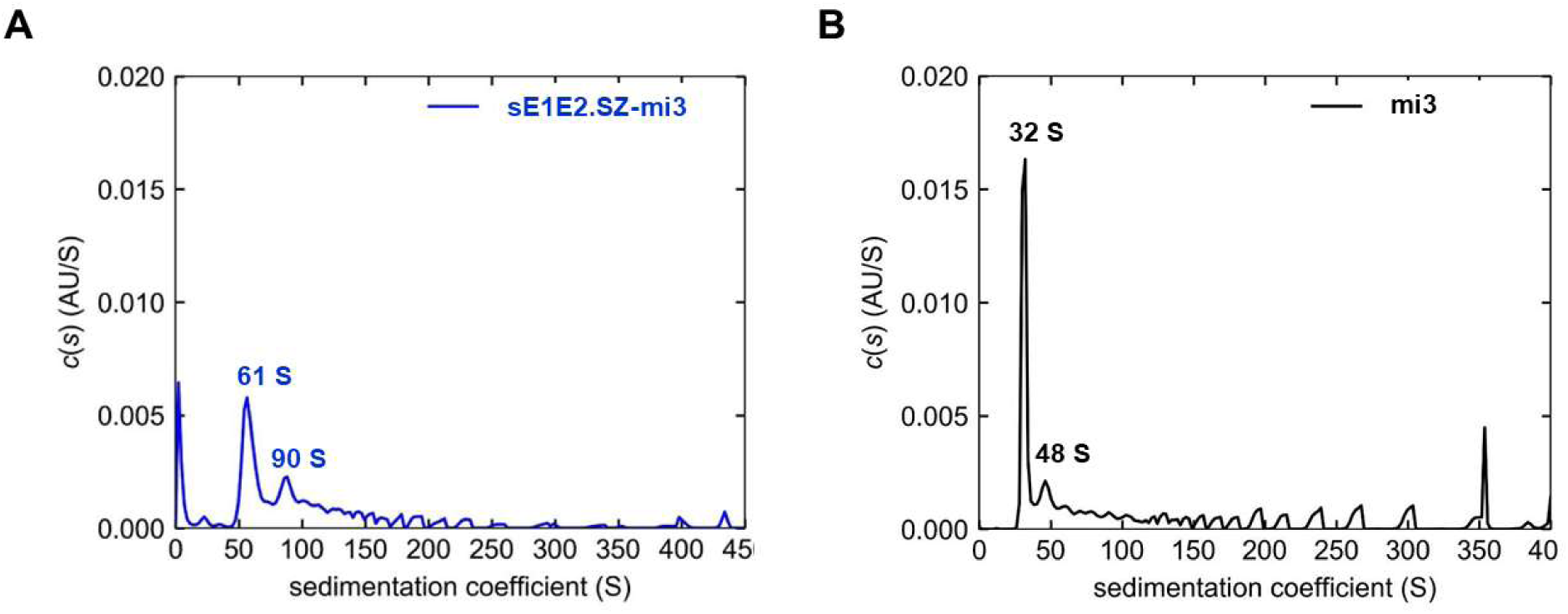
Analytical characterization of the size and heterogeneity of the sE1E2.SZ-mi3 and mi3 nanoparticles. AUC profiles of (A) sE1E2.SZ-mi3, and (B) mi3. Shown are the distribution of Lamm equation solutions c(s) for the two proteins (solid lines). Calculated sedimentation coefficients for the peaks are labeled. Both antigens have two major species. For sE1E2.SZ-mi3, the major species sediments at approximately 61 S, and the minor species sediments at approximately 90 S. For mi3, the major species sediments at approximately 32 S, and the minor species sediments at approximately 48 S.

### Characterization of antigenic integrity

We assessed the antigenic integrity of the sE1E2.SZ-mi3 nanoparticles by measuring binding to a panel of mAbs in comparison to sE1E2.SZ and mbE1E2. Dissociation constants (K_d_s) were measured by dose-dependent ELISA to antibodies that recognize discrete epitopes of E1, E2, and E1E2 (**Table 1**). The majority of these antibodies are conformation sensitive and span the commonly assessed antigenic domains on E2 (domains A, B, D, and E) and the metastable domains AR4 and AR5, both of which reside exclusively on E2 but require the presence of E1 to adopt their binding-competent conformations (25, 46, 47). For the majority of the antibodies, the differences in both K_d_ and maximum amplitude of the binding reaction (B_max_) are negligible.

**Table 1.** Affinity of antigens for mAbs. * Shown are the average and standard deviation of duplicate measurements

| Antibody | Domain | K<sub>d</sub> (nM)* |  |  |
| --- | --- | --- | --- | --- |
|  |  | sE1E2.SZ-mi3 | sE1E2.SZ | mbE1E2 |
| H-111 | E1 | 4.0 ± 0.1 | 0.40 ± 0.01 | 2.8 ± 0.9 |
| IGH526 | E1 | 1.6 ± 0.1 | 2.4 ± 0.1 | 6.3 ± 0.1 |
| CBH-4B | A | 25 ± 1 | 18 ± 1 | 22 ± 5 |
| AR3A | B | 0.39 ± 0.04 | 0.15 ± 0.01 | 0.13 ± 0.04 |
| HEPC74 | B | 0.33 ± 0.02 | 0.14 ± 0.01 | 0.22 ± 0.01 |
| HC84.26 | D | 0.17 ± 0.01 | 0.08 ± 0.01 | 0.14 ± 0.01 |
| HC84.1 | D | 0.22 ± 0.01 | 0.09 ± 0.01 | 0.16 ± 0.01 |
| HC33.1 | E | 0.07 ± 0.01 | 0.06 ± 0.01 | 2.2 ± 0.1 |
| HCV1 | E | 0.09 ± 0.01 | 0.07 ± 0.01 | 1.2 ± 0.1 |
| AR4A | AR4 | 0.83 ± 0.04 | 0.18 ± 0.01 | 0.15 ± 0.01 |
| AR5A | AR5 | 3.5 ± 0.1 | 0.76 ± 0.01 | 0.37 ± 0.06 |

There are, however, some differences in affinity. Both mbE1E2 and the sE1E2.SZ-mi3 nanoparticle bind with reduced affinity to the weakly-neutralizing mAb H-111, which binds to the E1 N-terminus, in comparison to sE1E2.SZ. The difference in affinity is approximately 10-fold, with a concomitant decrease in B_max_. Because the epitope recognized by the H-111 mAb participates in the E1E2 interface, it is possible that the multimeric nature of the mbE1E2 agglomerates (24) and the sE1E2.SZ-mi3 nanoparticles shield this epitope in some manner relative to the sE1E2.SZ heterodimer. Similarly, as we have observed previously (24), mbE1E2 binding to domain E antibodies HCV1 and HC33.1 is suppressed relative to both sE1E2.SZ and sE1E2.SZ-mi3. Finally, sE1E2.SZ-mi3 binding to the E1E2-specific mAbs AR4A and AR5A is decreased 5- to 9-fold relative to sE1E2.SZ and mbE1E2. In aggregate, these data indicate that the procedure we adopted for coupling sE1E2 to the mi3 nanoparticle scaffold retains the antigenic integrity of native E1E2. Surprisingly, we did not see an avidity effect for either the nanoparticles or mbE1E2 despite their multimeric quaternary structures. The reasons for this are not clear.

### Evaluation of the anti-E1E2 immune response by ELISA

Purified vaccine antigens were formulated into nanoscale size supramolecular assemblies with a polyphosphazene adjuvant co-formulated with the TLR7/8 agonist resiquimod (PCPP-R) as described previously (68–70). For the sE1E2.SZ-mi3 nanoparticle and sE1E2.SZ antigens, CD1 mice (n = 6 per group) were immunized using two separate concentration regimes (i.e. high and low), whereas for the mbE1E2 antigen and the control mi3 nanoparticle, mice were immunized at a single high concentration. Microgram amounts for the immunizations were adjusted based on the components of the antigen (e.g., sE1E2.SZ-mi3 contains sE1E2 and the nanoparticle shell) such that the equivalent amount of E1E2 was included in each immunization for the corresponding high and low concentration groups. Blood samples were collected prior to each vaccination on days 0 (pre-bleed), 14, 28, and 42, with a terminal bleed on day 56. At the conclusion of the study, pooled sera from each group were used to examine the kinetics of the anti-HCV E1E2 antibody response by assessing the overall titer at each collection point. We tested two different immobilized antigens, mbE1E2 and sE1E2.LZ (24) to evaluate the endpoint titer kinetics to assess potential differences in the immune response due to the truncations required to liberate sE1E2 from the membrane. We immobilized sE1E2.LZ instead of sE1E2.SZ to avoid confounding effects that might be caused by antibody binding to the exposed SZ scaffold. As shown in **Fig. 5A**, when sE1E2.LZ is the immobilized antigen, the sE1E2.SZ-mi3 high concentration group elicited an anti-E1E2 response (endpoint titer > 1000) at the first collection point after the initial immunization (day 14), whereas the titer for all other groups was at least 10-fold lower. Similarly, when mbE1E2 is the immobilized antigen, only the sera from the mbE1E2- and sE1E2.SZ-mi3-immunized mice contained a measurable endpoint titer (>100). By day 28, the endpoint titers for all groups had increased significantly (>25,000) for all groups tested against immobilized sE1E2.LZ except for the mi3 control, with the sE1E2.SZ-mi3 immunized mouse sera retaining the highest titer at this time point. When mbE1E2 is the immobilized antigen, the sera from mbE1E2-immunized mice exhibited a marked increase in titer (>50,000) as did the sE1E2.SZ-mi3 high concentration sera (>10,000), whereas the other groups had titers less than 5,000. By day 42, the titers had begun to plateau for all groups, with modest increases in titer for the groups at day 56. The mbE1E2-immunized sera retained a significantly higher titer than the sE1E2.SZ-mi3 and sE1E2.SZ groups when assessed against immobilized mbE1E2 (**Fig. 5B**). In contrast, all sera had similarly high titers at days 42 and 56 when sE1E2.LZ was the immobilized antigen. We then assessed the individual mouse endpoint titers at day 56 using the same two immobilized antigens (**Figs. 5C and D**). Consistent with the pooled sera results, when mbE1E2 is the immobilized antigen the mbE1E2-immunized mice exhibit a higher titer than the other groups and the endpoint titers all cluster around a similar mean when sE1E2.LZ is the immobilized antigen. The mbE1E2 mean endpoint titer is slightly lower at day 56 when sE1E2.LZ is the immobilized antigen, but this difference is not statistically significant. The spread in endpoint titer values also varies only modestly (from 2- to 6-fold) regardless of which antigen is immobilized on the plate. These results are consistent with previous observations using our first-generation scaffolded sE1E2 containing the c-Fos/c-Jun heterodimeric coiled coil scaffold (sE1E2.LZ) and comparing endpoint titers between sE1E2.LZ-immunized and mbE1E2-immunized mice (48). In those studies, we further showed that, relative to the mbE1E2-immunized mice, the sE1E2.LZ-immunized mice elicited fewer antibodies directed at hypervariable region 1 (HVR1) and antigenic domain A, both of which elicit primarily non-neutralizing antibodies (48).

**Figure 5.**
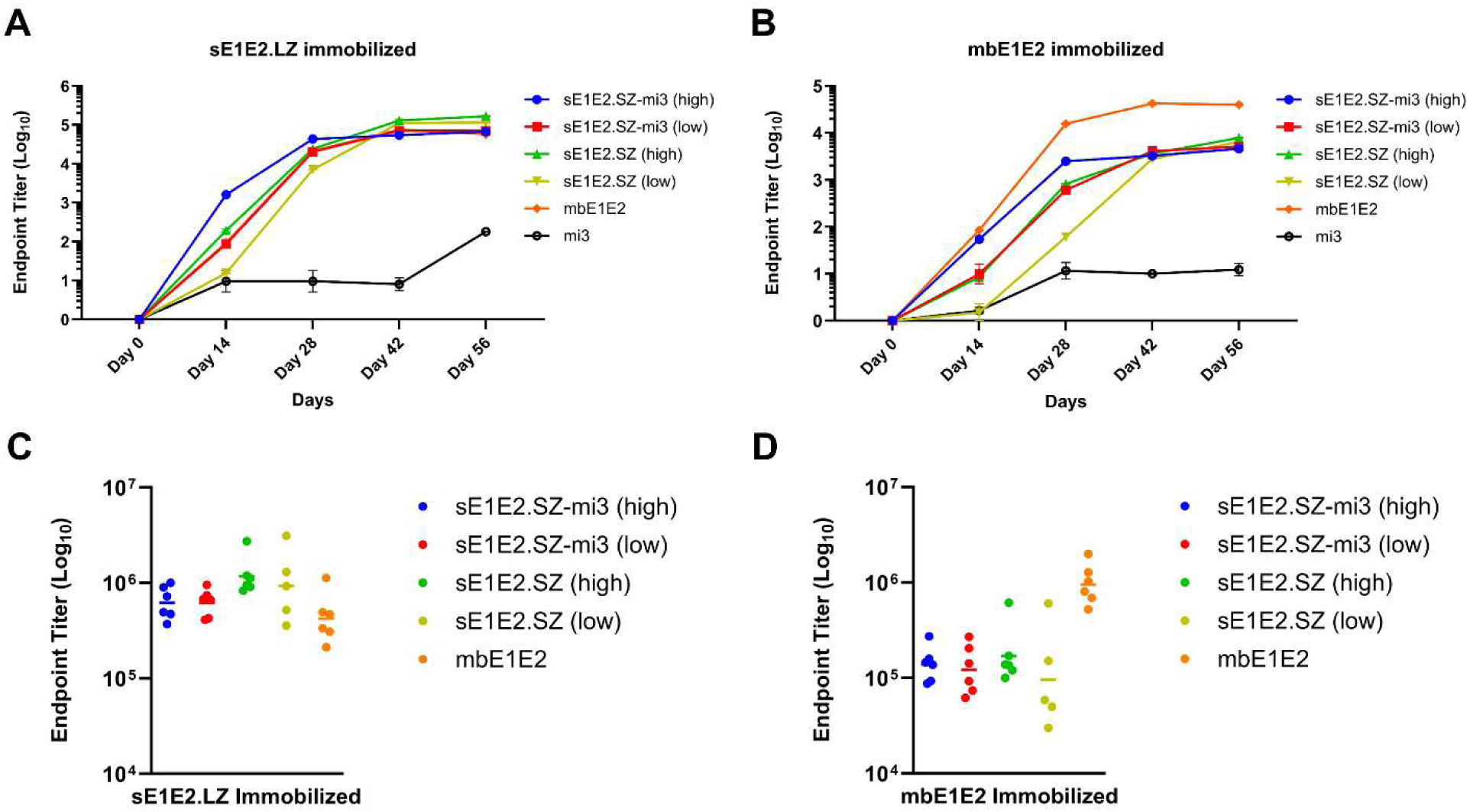
Assessment of antibodies induced in mice immunized with sE1E2.SZ-mi3 and mi3 by ELISA. Each datum represents the mean of duplicate experiments for each serum sample. **A.** Anti-sE1E2.LZ titers at different days post-immunization for each immunization group. **B.** Anti-mbE1E2 titers at different days post-immunization for each immunization group. **C.** Endpoint titers against the immobilized sE1E2.LZ antigen using individual mouse sera from day 56 post immunization. **D.** Endpoint titers against the immobilized mbE1E2 antigen using individual mouse sera from day 56 post immunization. Endpoint titers were calculated by curve fitting in GraphPad Prism software with the endpoint OD defined as four times the highest absorbance value of day 0 sera.

### Evaluation of serum bnAb response

The ability of sera from mice immunized with sE1E2.SZ-mi3 nanoparticles, sE1E2.SZ, and mbE1E2 to inhibit HCV infection *in vitro* was tested first against HCVpp containing the structural proteins from the homologous genotype 1a (H77) virus. Serum neutralization activities were first assessed using IgGs purified from pooled sera collected at days 0, 14, 28, 42, and 56. The kinetics of the bnAb response against the H77 HCVpp are shown in **Fig. 6** and **Fig. S2**. At day 14, which follows the initial prime immunization, the bnAb response was negligible, with pooled purified IgGs from only the two sE1E2.SZ-mi3 nanoparticle and mbE1E2 sera exhibiting neutralizing potential that exceeds that of the background day 0 IgGs. Purified IgGs from day 28, which follows the first boost immunization, exhibit a marked increase in H77 HCVpp neutralization, with both the high concentration sE1E2.SZ-mi3 group and the mbE1E2 group neutralizing greater than 90% of the HCVpp at the tested concentration (85 μg/mL). The low concentration sE1E2.SZ-mi3 pooled IgGs neutralize approximately 58% of the H77 HCVpp at this concentration, and pooled purified IgGs from the two sE1E2.SZ groups neutralize comparably less of the H77 HCVpp (approximately 48 and 39 % respectively). This trend is corroborated by examining the IC_50_ values for the pooled IgGs from each group. The mbE1E2 group exhibited the lowest IC_50_ (4.1 μg/mL), followed by the high concentration sE1E2.SZ-mi3 group (28.5 μg/mL), with both of the sE1E2.SZ groups exhibiting IC_50_ values in excess of 100 μg/mL (**Fig S2**). Thus, the response elicited by the sE1E2.SZ-mi3 nanoparticles exhibits an increase in neutralization potency relative to the sE1E2.SZ subunit vaccine after a prime and one boost such that it is nearly on par with that elicited by mbE1E2. By day 42, the bnAb response began to plateau, with pooled purified IgGs from all groups neutralizing in excess of 90% of the H77 HCVpp at the tested concentration. We then assessed heterologous neutralization at day 28, which is the time at which the disparity between homologous neutralization by the IgGs elicited by the antigens is greatest. Due to the background levels of neutralization exhibited by the IgGs from mice immunized with the antigens at low concentrations, we assessed only three groups: sE1E2.SZ-mi3 (high), sE1E2.SZ (high), and mbE1E2. As shown in **Fig 6B**, the subunit vaccine exhibited background levels of neutralization (<20%) for all heterologous pseudoviruses tested. By contrast, the IgGs from mice immunized with sE1E2-mi3 and mbE1E2 showed modest neutralization of 1b09 psuedovirus, but background levels of neutralization for J6, UKNP2.5.1, and UKNP3.2.2 pseudoviruses.

**Figure 6.**
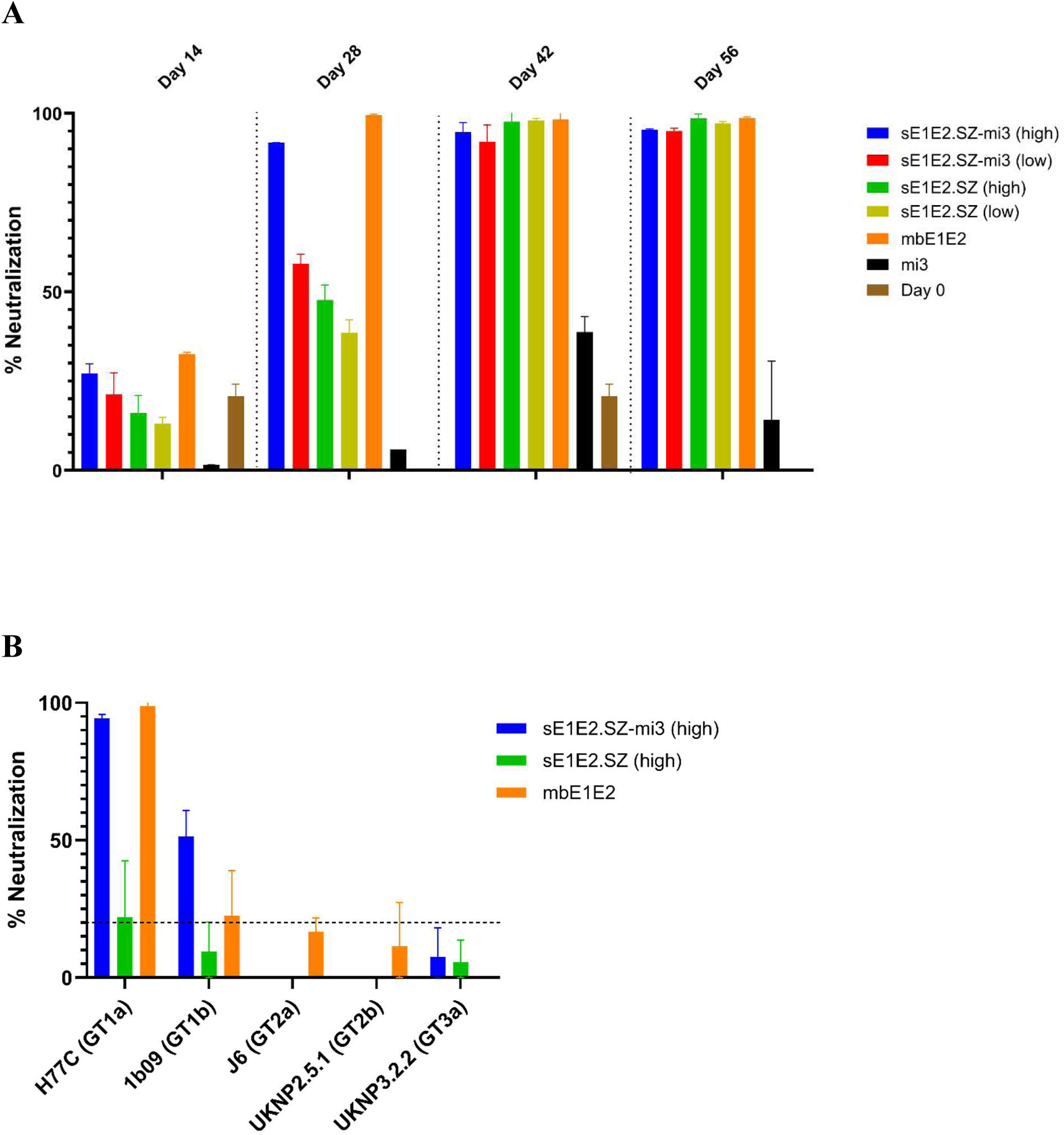
(A) Kinetics of homologous (H77C) pseudovirus neutralization by purified IgG from immunized mice. (B) Heterologous neutralization by purified IgG from immunized mice. A replicate of the homologous neutralization experiment from (A) is included as a positive control. The approximate level of background neutralization at day 28 is indicated by a dashed line. Each datum represents the mean of duplicate experiments for each IgG sample. IgGs were purified from pooled sera for each group at the indicated day post immunization and diluted to a final concentration of 85 µg/mL for neutralization experiments. Percent neutralization was calculated using relative luminescence units (RLU) normalized to RLU of supernatant cultured without HCVpp nor IgGs (100%) and RLU of supernatant cultured with HCVpp without IgGs (0%).

## Discussion

One factor that influences immunogenicity is size, which depends on the presentation mode of the vaccine antigen and affects antigen uptake and clearance. Subunit vaccines are typically monomeric or low-order oligomers (e.g. HIV env trimers (2)), whereas live and attenuated viruses present the vaccine antigen as polyvalent antigen complexes on the virion. For example, an inactivated form of SARS-CoV-2 is used as a vaccine (49), and the virus contains on average 26+/-15 spike trimers on its surface (50), whereas the purified spike subunit ectodomain is a trimer. One way to recapture some of the immunogenic potency of the virus vaccine system is to use a particulate or emulsion-based formulation (4, 5, 7). One common particulate formulation is Alum, which forms aggregated microparticles ranging from 0.5 to 10 µm in size (51). Other options include polymer-based nanoparticulates, such as poly(D,L-lactic-co-glycolic acid) or PLG (8, 52) and polyphosphazene (PPZ) (53–56) and micelle-forming pluronics (57), emulsions, such as MF59 (58) and Montanide (59), and nanovesicles, which include liposomes (60) - a basis for the advanced AS01 (61) and CAF01 (62) systems. For some membrane-associated glycoprotein vaccine candidates, the formulation process creates rosettes of antigen protruding from a hydrophobic micelle core, thereby creating nanoparticulate vaccines as in the cases of Flublok (63, 64) and the protein-based SARS-CoV-2 vaccine (65). Another means to increase the size of vaccine antigens is via incorporation on the surface of protein-based nanoparticles. Nanoparticle platforms for the multivalent display of vaccine antigens have proven effective in boosting immunogenicity for antigens such as Ebola GP (66), HIV env (67–69), SARS CoV-2 spike (70), RSV F (71), and influenza hemagglutinin (42). One method for constructing a protein-based nanoparticle vaccine is via direct genetic fusion. We recently participated in a collaborative effort which successfully produced an in cis genetic fusion HCV E1E2 NP vaccine candidate (23). The in cis NPs exhibited similar early induction of bNAbs that we observe here for the in trans HCV E1E2 NPs (23). In this study, we chose a method that leverages isopeptide bond formation to allow coupling of purified antigen to a purified nanoparticle scaffold. This “plug and display” modular assembly (17) uses the components SpyTag and SpyCatcher (45, 72) to create a covalent tethering system incorporated on the surface of the nanoparticles. The plug-and-display format is particularly useful for antigens that are difficult to produce in NP form as genetic fusions and/or for producing mosaic NPs in which a number of variants of a particular antigen are coupled simultaneously to the same NP in order to increase the breadth of neutralization of the vaccine (39, 42). This could be particularly useful against a pathogen like HCV, which is known to have a high degree of genetic diversity. Moreover, being able to approach nanoparticle assembly using both cis (23) and trans platforms increase the chances of success. We view the two approaches to be complementary, where the in cis platform could be a more efficient means of producing a particular strain or strains and the in trans platform could be more amenable to the mosaic vaccine approach.

In this study, we established a proof-of-principle method for assembling and E1E2-based nanoparticle vaccine using the plug-and-display system (17). The purified sE1E2.SZ was coupled to SpyCatcher-mi3 with high efficiency, as evidenced by western blotting and negative stain EM showing no unoccupied nanoparticles. The size of the nanoparticles themselves affords facile purification from uncoupled antigens via gel filtration chromatography. Based on our ELISA results, these nanoparticles are antigenically intact post-coupling and purification. The sE1E2.SZ-mi3 group showed an anti-E1E2 response beginning at day 14, i.e., after the priming dose. Moreover, sE1E2.SZ-mi3 elicited a bnAb response against the homologous strain starting as early as day 28 that was comparable to the response elicited by mbE1E2, which is known to form large agglomerates in solution (38). In contrast, the bnAb response against the homologous strain was weaker for the subunit vaccine groups. Moreover, purified IgGs from the sE1E2.SZ-mi3-immunized mice exhibited cross-neutralizing activity against the 1b09 strain at day 28. This neutralizing activity is more potent than the neutralizing activity of the IgGs from mice immunized with mbE1E2 against 1b09. The IgGs from mice immunized with the sE1E2.SZ subunit vaccine exhibited only background levels of neutralization against the 1b09 strain. The 1b09 strain is classified as a tier 2 strain and has a resistance to neutralization similar to that of H77 (73), whereas the other heterologous strains are more resistant (74, 75). Thus, the increased valency of sE1E2.SZ-mi3 relative to its subunit vaccine appears to enhance the bnAb response after a prime dose and a single boost. Minimization of the number of immunizations required to elicit a bnAb response will be an important factor in developing an effective HCV vaccine as one of the major cohorts at high risk of HCV infection and thus would benefit most from such a vaccine would be people who inject drugs (76). Based on the above data, these studies represent a proof-of-principle demonstration that the well-characterized sE1E2.SZ platform compatible with multiple genotypes can be coupled to a nanoparticle system that is currently in use in clinical trials (44, 77).

## Materials and Methods

### Plasmid construction

The clone for expressing membrane-bound E1E2 (mbE1E2) has been described previously (24). The soluble secreted sE1E2.SZ.SpyTag clone was based on the previously described sE1E2.SZ construct (25) and modified to incorporate an optimized version of the SpyTag (45) at the C-terminus of the SYNZIP1 scaffold appended to the E2 ectodomain. For simplicity, we will refer to this as sE1E2.SZ throughout the manuscript. The optimized SpyCatcher-mi3 (45) open reading frame was synthesized by GenScript and subcloned into pET17b for bacterial expression.

### Protein expression and purification

Expression of recombinant sE1E2.SZ and mbE1E2 was performed via transient expression in human Expi293 cells using the Expi293 Expression System by following the manufacturer’s protocols (Thermo Fisher Scientific, Walthan, MA). Briefly, Expi293 cells were cultured in Expi293 Expression Medium in the shaker incubator at 37 °C, with 120 rpm and 8% CO_2_. When the cells reached a density of 2.0 × 10^6^ cells/mL, Expi293 cells were transfected using proper amounts of plasmid DNA. Culture supernatants of sE1E2.SZ were harvested at 72 hours after transfection, clarified by centrifugation at 10,000 rpm for 10 min, and filtered by a 0.22 μm filters. sE1E2.SZ was purified using our established protocol, which entailed subjecting the clarified supernatant to sequential HiTrap chelating sepaharose and Superdex 200 size exclusion chromatography (SEC) as described in our previous papers (24, 78). Expi293 cells transfected with recombinant mbE1E2 were collected 72 hours after transfection and the cell pellets were lysed using 1% NP-9 cell lysis buffer (24). Recombinant mbE1E2 was then purified by sequential Fractogel EMD TMAE (Millipore), Fractogel EMD SO3- (Millipore), and HC84.26 (79) immunoaffinity chromatography as described previously (24, 48).

### SDS-PAGE and western blot

Purified sE2 antigens were separated by a precast, 4–20% Mini-PROTEAN TGX stain-free gels on a Mini-PROTEAN Tetra cell electrophoresis instrument (Bio-Rad Laboratories, Hercules, CA). In reducing conditions, each sample was incubated with loading dye (4x Laemmli buffer + 10% β-mercaptoethanol) (Bio-Rad Laboratories, Hercules, CA) and heated to 95 °C. In non-reducing conditions, each sample was incubated with Laemmli buffer and heated to 37 °C. For western blot detection, the purified protein samples on SDS-PAGE were transferred onto Trans-Blot Turbo Mini nitrocellulose membranes (Bio-Rad Laboratories, Hercules, CA). The membranes were then probed using the anti-HCV E2 mAb HCV1 (80) at 5 μg/mL or anti-HCV E1 mAb H111 (81) at 10 μg/mL followed by detection using a secondary goat anti-human IgG-HRP conjugate (Invitrogen, Waltham, MA) at a 0.16 μg/mL dilution and the Western ECL substrate (Bio-Rad Laboratories, Hercules, CA). All gels were imaged using the ChemiDoc system (Bio-Rad Laboratories, Hercules, CA).

### Negative stain electron microscopy

Preliminary negative stain EM studies were conducted at the Keith R. Porter Imaging Facility at the University of Maryland, Baltimore County (UMBC). For these samples, 10 µL of sample was placed on to 200 mesh formvar-covered, carbon-coated copper grids (EMS, Hatfield, PA, USA), incubated for 1 min, briefly rinsed with ultra-pure water, then stained with 2% uranyl acetate for 2 min. Images were taken at 100 kV using a Hitachi HT7800 120 kV TEM equipped with an AMT Nanosprint15 B digital camera. Analysis of these initial images was performed using cisTEM (82).

For samples prepared at larger scale for animal studies, TEM grids were prepared following general protocol described in Rames, et al. (83). Briefly, 4 µL of a sample were deposited on a glow discharged continuous carbon film 300 mesh copper grids (Electron Microscopy Science, Hatfield, PA) and incubated for 1 min to allow for the particle absorption on the carbon film. Excess liquid was then removed by gently touching the side of the grid with a filter paper. This was followed by washing the grid 3 times with deionized water by touching a fresh water drop and instantly removing excess liquid. This process was then repeated with 3 fresh drops of 1% uranyl formate, with short incubations at the first two steps (∼10 seconds) followed by a longer 1 minute staining at the last step. Grids were then allowed to dry overnight at room temperature.

TEM data were collected on FEI Talos Arctica 200kV electron microscope equipped with Gatan K3 direct electron detector. Since the microscope lacks an anticontamination device, data collection was performed at cryogenic temperatures using frozen grids. Several hundred micrographs were collected for each sample at low dose conditions at ∼56,000x magnification using SerialEM software. Cryo-EM movies were patch motion corrected using cryoSPARC v2.15(84). CTF parameters were estimated using the Patch CTF job within cryoSPARC. Blob picking was implemented on a subset of the dataset, and the resulting particles were used for 2D classification. Particles corresponding to 2D classes of the corresponding nanoparticles were used for *ab initio* reconstruction. After selecting 2D classes consisting of the desired nanoparticle multiple rounds of 3D classification were performed. For the sE1E2-mi3 3D classification, no symmetry was imposed and for the mi3 alone 3D classification, isocahedral symmetry was imposed to achieve the final reconstruction.

### Analytical Ultracentrifugation

Sedimentation velocity (SV) experiments were performed at 20 °C using a ProteomeLab Beckman XL-A with absorbance optical system and a 4-hole An60-Ti rotor (Beckman Coulter, Indianapolis, IN). For both the sE1E2.SZ-mi3 nanoparticles and the control mi3 nanoparticles, the sample and reference sectors of the dual-sector charcoal-filled epon centerpieces were loaded with 380 μL protein in PBS, pH 7.4, and 400 μL buffer. The cells were centrifuged at 20 krpm and the absorbance data were collected at 280 nm in a continuous mode with a step size of 0.003 cm and a single reading per step to obtain linear signals of <1.25 absorbance units.

Sedimentation coefficients were calculated from SV profiles using the program SEDFIT (85). The continuous *c*(*s*) distributions were calculated assuming a direct sedimentation boundary model with maximum entry regularization at a confidence level of 1 standard deviation. The density and viscosity of buffers at 20 °C were calculated using SEDNTERP (86). The *c*(*s*) distribution profiles were prepared with the program GUSSI (C.A. Brautigam, Univ. of Texas Southwestern Medical Center).

### ELISA screening of antigens for mAb a binding

HCV HMAb binding to sE1E2.SZ-mi3 nanoparticles, sE1E2.SZ, and mbE1E2 were evaluated and quantitated by ELISA. 96-well microplates (MaxiSorp, Thermo Fisher Scientific, Waltham, MA) were coated with 5 μg/mL Galanthus Nivalis Lectin (Vector Laboratories, Newark, CA) overnight, and purified antigen was then added to the plates at 2 μg/mL. After the plates were washed with PBS and 0.05% Tween 20, and blocked by Pierce™ Protein-Free (PBS) Blocking Buffer (Thermo Fisher Scientific, Waltham, MA), the mAbs were tested in duplicate at 3-fold serial dilution starting at a concentration of 66 nM. The binding was detected by HRP-conjugated anti-human IgG secondary antibody (Invitrogen, Waltham, MA) at a concentration of 0.16 mg/mL with TMB substrate (Bio-Rad Laboratories, Hercules, CA). The absorbance was read at 450 nm using a SpectraMax MS microplate reader (Molecular Devices, San Jose, CA). The data were analyzed by nonlinear regression to measure antibody dissociation constants (K_d_) using GraphPad Prism software.

### Animal immunization

CD1 mice were purchased from Charles River Laboratories. Prior to immunization, purified antigens were formulated with polyphosphazene adjuvant as described in previous studies (87, 88). In brief, 50 μg poly[di(carboxylatophenoxy)phosphazene (PCPP) was formulated with 25 μg resiquimod, R848 in PBS (pH 7.4) to form the PCPP-R adjuvant. The resulting supramolecular complex (PCPP-R) was formulated with each antigen. The dose was calibrated based on the composition of the antigen to ensure that an equal amount of E1E2 was administered for the corresponding high and low concentration groups. Thus, for the high concentration sE1E2.SZ-mi3 group, 100 μg was used for prime and 20 μg for boost immunization. For the low concentration sE1E2.SZ-mi3 group, 20 μg was used for prime and 4 μg for boost immunization. For the high concentration sE1E2.SZ group, 70 μg was used for prime and 15 μg for boost immunization. For the low concentration sE1E2.SZ group, 14 μg was used for prime and 3 μg for boost immunization. For the mbE1E2 group, 70 μg was used for prime and 15 μg for boost immunization. For the SpyTag-mi3 control group, 30 μg was used for prime and 6 μg for boost immunization. This approximates the amount of mi3 contained in the sE1E2.SZ-mi3 high concentration group. Dynamic light scattering (DLS) was used to confirm the absence of aggregation in adjuvanted formulations. Groups of six female CD-1 mice, age 7 to 9 weeks, were immunized via the intraperitoneal (IP) route, first with a prime as described above on day 0, then with boosts as described above on day 14, day 28 and day 42. Blood samples were collected prior to each vaccination on days 0 (pre-bleed), 14, 28, 42 and a terminal bleeding on day 56. The blood samples were processed for serum by centrifugation and stored at −80 °C until analysis was performed.

### ELISAs for serum antibody detection

ELISA was performed to measure HCV E1E2-specific antibody responses in sera from immunized mice. 96-well plates (MaxiSorp, Thermo Fisher Scientific, Waltham, MA) were coated overnight with 5 µg/mL Galanthus Nivalis Lectin (Vector Laboratories, Newark, CA) at 4°C. The next day, plates were washed with PBS containing 0.05% Tween 20 and coated with 200 ng/well of either sE1E2.LZ (24) or mbE1E2 at 4 °C. After overnight incubation, plates were washed with PBS containing 0.05% Tween 20 and blocked with Pierce™ Protein-Free Blocking Buffer (Thermo Fisher Scientific, Waltham, MA) for 1 hour, and serially diluted mice sera samples were then added to the plates and incubated for another hour. The binding of HCV E1E2-specific antibodies was detected by an HRP-conjugated anti-mouse IgG secondary antibody (Abcam, Waltham, MA) at a concentration of 0.4 μg/mL with TMB substrates (Bio-Rad Laboratories, Hercules, CA). Absorbance values at 450 nm (SpectraMax M3 microplate reader) were used to determine endpoint titers, which were calculated by curve fitting in GraphPad Prism software and defined as four times the highest absorbance value of pre-immune sera. Statistical analysis to determine significance was performed using a Kruskal-Wallis one-way ANOVA.

### HCVpp neutralization assay

Total serum IgGs were purified using protein G HP SpinTrap columns (Cytiva, Marlborough, MA) according to the manufacturer’s protocol. Briefly, 600 μL of mouse serum was loaded on the column and incubated for 4 min with gentle mixing. After washing 2 times with binding buffer (20 mM sodium phosphate, pH 7.0), IgGs were eluted with elution buffer (0.1 M glycine-HCl, pH 2.7) into tubes containing neutralizing buffer (1 M Tris-HCl, pH 9.0). For HCVpp neutralization, purified IgGs were serially diluted from a starting concentration of 100 μg/mL.

The human hepatoma cell line, Huh7, was maintained in the DMEM medium supplemented with 10 % FBS and 1% non-essential amino acids (NEAA) (Thermo Fisher Scientific, Waltham, MA), and used as the target cell line for neutralization assays (24, 89). To test antibodies and purified IgG for neutralization, Huh7 cells were pre-seeded into 96-wells plates at a density of 1 × 10^4^ per well. In next day, the pseudoparticles were incubated with defined concentrations of mAbs and/or the purified IgGs at indicated concentrations for 1 hour at 37 °C, and then added to each well. After the plates were incubated in a CO_2_ incubator at 37 °C for 5 to 6 hours, the mixtures were replaced with fresh medium and then continued to incubate for 72 hours. After incubation, 100 μL of Bright-Glo reagent (Promega, Madison, WI) was added to each well for 2 minutes at room temperature and the luciferase activity was measured using a FLUOstar Omega plate reader (BMG Labtech, Cary, NC) with the MARS software. The 50% inhibitory concentration (IC_50_) was calculated as the mAb concentration that caused a 50% reduction in relative light units (RLU) compared with pseudoparticles in the control wells. All values were calculated using a dose-response curve fit with nonlinear regression in GraphPad Prism. All experiments involving the use of pseudoparticles were performed under biosafety level 2 conditions. Statistical analyses to determine significance were performed using Kruskal–Wallis analysis of variance with Dunn’s multiple comparison test.

## ACKNOWLEDGEMENTS

This work was supported by NIAID (R01 AI168048, T.R.F., E.A.T., A.A., B.G.P.) and by the Maryland Center for Advanced Molecular Analyses and the MPowering the State program in Maryland (E.P. and T.R.F.).

## DATA AVAILABILITY

The data supporting the results of this study are included in the article/supplementary material. Further inquiries can be directed to the corresponding authors.

## AUTHOR CONTRIBUTIONS

T.R.F., E.A.T., and B.G.P. developed the concept of the project. T.R.F., E.A.T., B.G.P., K.L.C., T.C., E.P., Y.L., and A.K.A. designed the experiments. L.K., S.J., D.Z, K.L.C., M.C.M, A.M., X.S., T.C., and E.P. conducted experiments. L.K., S.J., D.Z., K.L.C., T.C., E.P. A.M., M.C.M., and E.A.T. performed data analysis. L.K., T.R.F., B.G.P., T.C., E.P., Y.L., and E.A.T. wrote the manuscript. T.R.F coordinated the animal study. All authors approve of the submitted manuscript.

## REFERENCES

1. Sookhoo JRV, Schiffman Z, Ambagala A, Kobasa D, Pardee K, Babiuk S. 2024. Protein Expression Platforms and the Challenges of Viral Antigen Production. Vaccines (Basel) 12.

2. Center RJ, Leapman RD, Lebowitz J, Arthur LO, Earl PL, Moss B. 2002. Oligomeric structure of the human immunodeficiency virus type 1 envelope protein on the virion surface. J Virol 76:7863–7867.

3. Akazawa D, Moriyama M, Yokokawa H, Omi N, Watanabe N, Date T, Morikawa K, Aizaki H, Ishii K, Kato T, Mochizuki H, Nakamura N, Wakita T. 2013. Neutralizing antibodies induced by cell culture-derived hepatitis C virus protect against infection in mice. Gastroenterology 145:447–455 e441-444.

4. Al-Halifa S, Gauthier L, Arpin D, Bourgault S, Archambault D. 2019. Nanoparticle-Based Vaccines Against Respiratory Viruses. Front Immunol 10:22.

5. Kim D, Wu Y, Kim YB, Oh YK. 2021. Advances in vaccine delivery systems against viral infectious diseases. Drug Deliv Transl Res 11:1401–1419.

6. Kushnir N, Streatfield SJ, Yusibov V. 2012. Virus-like particles as a highly efficient vaccine platform: diversity of targets and production systems and advances in clinical development. Vaccine 31:58–83.

7. Zhou J, Kroll AV, Holay M, Fang RH, Zhang L. 2020. Biomimetic Nanotechnology toward Personalized Vaccines. Adv Mater 32:e1901255.

8. De Temmerman ML, Rejman J, Demeester J, Irvine DJ, Gander B, De Smedt SC. 2011. Particulate vaccines: on the quest for optimal delivery and immune response. Drug Discov Today 16:569–582.

9. Aung A, Cui A, Maiorino L, Amini AP, Gregory JR, Bukenya M, Zhang Y, Lee H, Cottrell CA, Morgan DM, Silva M, Suh H, Kirkpatrick JD, Amlashi P, Remba T, Froehle LM, Xiao S, Abraham W, Adams J, Love JC, Huyett P, Kwon DS, Hacohen N, Schief WR, Bhatia SN, Irvine DJ. 2023. Low protease activity in B cell follicles promotes retention of intact antigens after immunization. Science 379:eabn8934.

10. Bale JB, Gonen S, Liu Y, Sheffler W, Ellis D, Thomas C, Cascio D, Yeates TO, Gonen T, King NP, Baker D. 2016. Accurate design of megadalton-scale two-component icosahedral protein complexes. Science 353:389–394.

11. Bruun TUJ, Andersson AC, Draper SJ, Howarth M. 2018. Engineering a Rugged Nanoscaffold To Enhance Plug-and-Display Vaccination. ACS Nano 12:8855–8866.

12. Domingo GJ, Orru S, Perham RN. 2001. Multiple display of peptides and proteins on a macromolecular scaffold derived from a multienzyme complex. J Mol Biol 305:259–267.

13. Hsia Y, Bale JB, Gonen S, Shi D, Sheffler W, Fong KK, Nattermann U, Xu C, Huang PS, Ravichandran R, Yi S, Davis TN, Gonen T, King NP, Baker D. 2016. Design of a hyperstable 60-subunit protein dodecahedron. [corrected]. Nature 535:136–139.

14. Lopez-Sagaseta J, Malito E, Rappuoli R, Bottomley MJ. 2016. Self-assembling protein nanoparticles in the design of vaccines. Comput Struct Biotechnol J 14:58–68.

15. Wang W, Liu Z, Zhou X, Guo Z, Zhang J, Zhu P, Yao S, Zhu M. 2019. Ferritin nanoparticle-based SpyTag/SpyCatcher-enabled click vaccine for tumor immunotherapy. Nanomedicine 16:69–78.

16. Wei Y, Kumar P, Wahome N, Mantis NJ, Middaugh CR. 2018. Biomedical Applications of Lumazine Synthase. J Pharm Sci 107:2283–2296.

17. Brune KD, Leneghan DB, Brian IJ, Ishizuka AS, Bachmann MF, Draper SJ, Biswas S, Howarth M. 2016. Plug-and-Display: decoration of Virus-Like Particles via isopeptide bonds for modular immunization. Sci Rep 6:19234.

18. He L, Tzarum N, Lin X, Shapero B, Sou C, Mann CJ, Stano A, Zhang L, Nagy K, Giang E, Law M, Wilson IA, Zhu J. 2020. Proof of concept for rational design of hepatitis C virus E2 core nanoparticle vaccines. Sci Adv 6:eaaz6225.

19. Sliepen K, Radic L, Capella-Pujol J, Watanabe Y, Zon I, Chumbe A, Lee WH, de Gast M, Koopsen J, Koekkoek S, Del Moral-Sanchez I, Brouwer PJM, Ravichandran R, Ozorowski G, King NP, Ward AB, van Gils MJ, Crispin M, Schinkel J, Sanders RW. 2022. Induction of cross-neutralizing antibodies by a permuted hepatitis C virus glycoprotein nanoparticle vaccine candidate. Nat Commun 13:7271.

20. Yan Y, Wang X, Lou P, Hu Z, Qu P, Li D, Li Q, Xu Y, Niu J, He Y, Zhong J, Zhong H. 2020. A Nanoparticle-Based Hepatitis C Virus Vaccine With Enhanced Potency. **J Infect Dis** 221:1304**-**1314.

21. Kinchen VJ, Massaccesi G, Flyak AI, Mankowski MC, Colbert MD, Osburn WO, Ray SC, Cox AL, Crowe JE, Jr., Bailey JR. 2019. Plasma deconvolution identifies broadly neutralizing antibodies associated with hepatitis C virus clearance. J Clin Invest 129:4786–4796.

22. Frumento N, Figueroa A, Wang T, Zahid MN, Wang S, Massaccesi G, Stavrakis G, Crowe JE, Jr., Flyak AI, Ji H, Ray SC, Shaw GM, Cox AL, Bailey JR. 2022. Repeated exposure to heterologous hepatitis C viruses associates with enhanced neutralizing antibody breadth and potency. The Journal of Clinical Investigation 132.

23. He L, Lee Y-Z, Zhang Y-N, Newby ML, Janus BM, Gonzalez FG, Ward G, DesRoberts C, Hung S-H, Giang E, Allen JD, Kulakova L, Toth EA, Fuerst TR, Law M, Ofek G, Crispin M, Zhu J. 2026. Native-like soluble E1E2 glycoprotein heterodimers on self-assembling protein nanoparticles for hepatitis C virus vaccine design. Nature Communications doi:10.1038/s41467-026-69418-9.

24. Guest JD, Wang R, Elkholy KH, Chagas A, Chao KL, Cleveland TEt, Kim YC, Keck ZY, Marin A, Yunus AS, Mariuzza RA, Andrianov AK, Toth EA, Foung SKH, Pierce BG, Fuerst TR. 2021. Design of a native-like secreted form of the hepatitis C virus E1E2 heterodimer. Proc Natl Acad Sci U S A 118:e2015149118.

25. Metcalf MC, Janus BM, Yin R, Wang R, Guest JD, Pozharski E, Law M, Mariuzza RA, Toth EA, Pierce BG, Fuerst TR, Ofek G. 2023. Structure of engineered hepatitis C virus E1E2 ectodomain in complex with neutralizing antibodies. Nat Commun 14:3980.

26. Who. 2024. Global Hepatitis Report 2024. World Health Organization.

27. Anonymous. 2021. World Health Organization. Global progress report on HIV, viral hepatitis and sexually transmitted infections, 2021. https://www.who.int/publications/i/item/9789240027077. Accessed May 8, 2023.

28. Liang TJ, Ghany MG. 2013. Current and future therapies for hepatitis C virus infection. N Engl J Med 368:1907–1917.

29. Scheel TK, Rice CM. 2013. Understanding the hepatitis C virus life cycle paves the way for highly effective therapies. Nat Med 19:837–849.

30. Cox AL. 2015. MEDICINE. Global control of hepatitis C virus. Science 349:790–791.

31. Bowen DG, Walker CM. 2005. Adaptive immune responses in acute and chronic hepatitis C virus infection. Nature 436:946–952.

32. Mehta SH, Cox A, Hoover DR, Wang XH, Mao Q, Ray S, Strathdee SA, Vlahov D, Thomas DL. 2002. Protection against persistence of hepatitis C. Lancet 359:1478–1483.

33. Osburn WO, Fisher BE, Dowd KA, Urban G, Liu L, Ray SC, Thomas DL, Cox AL. 2010. Spontaneous control of primary hepatitis C virus infection and immunity against persistent reinfection. Gastroenterology 138:315–324.

34. Page K, Hahn JA, Evans J, Shiboski S, Lum P, Delwart E, Tobler L, Andrews W, Avanesyan L, Cooper S, Busch MP. 2009. Acute hepatitis C virus infection in young adult injection drug users: a prospective study of incident infection, resolution, and reinfection. J Infect Dis 200:1216–1226.

35. Honegger JR, Zhou Y, Walker CM. 2014. Will there be a vaccine to prevent HCV infection? Semin Liver Dis 34:79–88.

36. Man John Law L, Landi A, Magee WC, Lorne Tyrrell D, Houghton M. 2013. Progress towards a hepatitis C virus vaccine. Emerg Microbes Infect 2:e79.

37. Tarr AW, Khera T, Hueging K, Sheldon J, Steinmann E, Pietschmann T, Brown RJ. 2015. Genetic Diversity Underlying the Envelope Glycoproteins of Hepatitis C Virus: Structural and Functional Consequences and the Implications for Vaccine Design. Viruses 7:3995–4046.

38. Toth EA, Chagas A, Pierce BG, Fuerst TR. 2021. Structural and Biophysical Characterization of the HCV E1E2 Heterodimer for Vaccine Development. Viruses 13.

39. Cohen AA, Gnanapragasam PNP, Lee YE, Hoffman PR, Ou S, Kakutani LM, Keeffe JR, Wu HJ, Howarth M, West AP, Barnes CO, Nussenzweig MC, Bjorkman PJ. 2021. Mosaic nanoparticles elicit cross-reactive immune responses to zoonotic coronaviruses in mice. Science 371:735–741.

40. Cohen AA, van Doremalen N, Greaney AJ, Andersen H, Sharma A, Starr TN, Keeffe JR, Fan C, Schulz JE, Gnanapragasam PNP, Kakutani LM, West AP, Jr., Saturday G, Lee YE, Gao H, Jette CA, Lewis MG, Tan TK, Townsend AR, Bloom JD, Munster VJ, Bjorkman PJ. 2022. Mosaic RBD nanoparticles protect against challenge by diverse sarbecoviruses in animal models. Science 377:eabq0839.

41. Wang E, Cohen AA, Caldera LF, Keeffe JR, Rorick AV, Adia YM, Gnanapragasam PNP, Bjorkman PJ, Chakraborty AK. 2025. Designed mosaic nanoparticles enhance cross-reactive immune responses in mice. Cell 188:1036–1050 e1011.

42. Cohen AA, Yang Z, Gnanapragasam PNP, Ou S, Dam KA, Wang H, Bjorkman PJ. 2021. Construction, characterization, and immunization of nanoparticles that display a diverse array of influenza HA trimers. PLoS One 16:e0247963.

43. Kanekiyo M, Joyce MG, Gillespie RA, Gallagher JR, Andrews SF, Yassine HM, Wheatley AK, Fisher BE, Ambrozak DR, Creanga A, Leung K, Yang ES, Boyoglu-Barnum S, Georgiev IS, Tsybovsky Y, Prabhakaran MS, Andersen H, Kong WP, Baxa U, Zephir KL, Ledgerwood JE, Koup RA, Kwong PD, Harris AK, McDermott AB, Mascola JR, Graham BS. 2019. Mosaic nanoparticle display of diverse influenza virus hemagglutinins elicits broad B cell responses. Nat Immunol 20:362–372.

44. Yang RS, Traver M, Barefoot N, Stephens T, Alabanza C, Manzella-Lapeira J, Zou G, Wolff J, Li Y, Resto M, Shadrick W, Yang Y, Ivleva VB, Tsybovsky Y, Carlton K, Brzostowski J, Gall JG, Lei QP. 2024. Mosaic quadrivalent influenza vaccine single nanoparticle characterization. Sci Rep 14:4534.

45. Keeble AH, Turkki P, Stokes S, Khairil Anuar INA, Rahikainen R, Hytonen VP, Howarth M. 2019. Approaching infinite affinity through engineering of peptide-protein interaction. Proc Natl Acad Sci U S A 116:26523–26533.

46. Giang E, Dorner M, Prentoe JC, Dreux M, Evans MJ, Bukh J, Rice CM, Ploss A, Burton DR, Law M. 2012. Human broadly neutralizing antibodies to the envelope glycoprotein complex of hepatitis C virus. Proc Natl Acad Sci U S A 109:6205–6210.

47. Torrents de la Pena A, Sliepen K, Eshun-Wilson L, Newby ML, Allen JD, Zon I, Koekkoek S, Chumbe A, Crispin M, Schinkel J, Lander GC, Sanders RW, Ward AB. 2022. Structure of the hepatitis C virus E1E2 glycoprotein complex. Science 378:263–269.

48. Wang R, Suzuki S, Guest JD, Heller B, Almeda M, Andrianov AK, Marin A, Mariuzza RA, Keck ZY, Foung SKH, Yunus AS, Pierce BG, Toth EA, Ploss A, Fuerst TR. 2022. Induction of broadly neutralizing antibodies using a secreted form of the hepatitis C virus E1E2 heterodimer as a vaccine candidate. Proc Natl Acad Sci U S A 119:e2112008119.

49. Hotez PJ, Bottazzi ME. 2022. Whole Inactivated Virus and Protein-Based COVID-19 Vaccines. Annu Rev Med 73:55–64.

50. Yao H, Song Y, Chen Y, Wu N, Xu J, Sun C, Zhang J, Weng T, Zhang Z, Wu Z, Cheng L, Shi D, Lu X, Lei J, Crispin M, Shi Y, Li L, Li S. 2020. Molecular Architecture of the SARS-CoV-2 Virus. Cell 183:730–738 e713.

51. Orr MT, Khandhar AP, Seydoux E, Liang H, Gage E, Mikasa T, Beebe EL, Rintala ND, Persson KH, Ahniyaz A, Carter D, Reed SG, Fox CB. 2019. Reprogramming the adjuvant properties of aluminum oxyhydroxide with nanoparticle technology. npj Vaccines 4:1.

52. Jain S, O’Hagan DT, Singh M. 2011. The long-term potential of biodegradable poly(lactideco-glycolide) microparticles as the next-generation vaccine adjuvant. Expert Rev Vaccines 10:1731–1742.

53. Andrianov AK, Langer R. 2021. Polyphosphazene immunoadjuvants: Historical perspective and recent advances. Journal of Controlled Release 329:299–315.

54. Chand DJ, Magiri RB, Wilson HL, Mutwiri GK. 2021. Polyphosphazenes as adjuvants for animal vaccines and other medical applications. Frontiers in Bioengineering and Biotechnology 9:625482.

55. Eng NF, Garlapati S, Gerdts V, Potter A, Babiuk LA, Mutwiri GK. 2010. The potential of polyphosphazenes for delivery of vaccine antigens and immunotherapeutic agents. Current drug delivery 7:13–20.

56. Andrianov AK, Fuerst TR. 2021. Immunopotentiating and Delivery Systems for HCV Vaccines. Viruses 13.

57. Coeshott CM, Smithson SL, Verderber E, Samaniego A, Blonder JM, Rosenthal GJ, Westerink MAJ. 2004. Pluronic® F127-based systemic vaccine delivery systems. Vaccine 22:2396–2405.

58. O’Hagan DT, Ott GS, Nest GV, Rappuoli R, Giudice GD. 2013. The history of MF59? adjuvant: a phoenix that arose from the ashes. Expert Rev Vaccines 12:13–30.

59. van Doorn E, Liu H, Huckriede A, Hak E. 2016. Safety and tolerability evaluation of the use of Montanide ISA™51 as vaccine adjuvant: A systematic review. Hum Vaccines Immunother 12:159–169.

60. Schwendener RA. 2014. Liposomes as vaccine delivery systems: a review of the recent advances. Ther Adv Vaccines 2:159–182.

61. Didierlaurent AM, Laupèze B, Di Pasquale A, Hergli N, Collignon C, Garçon N. 2017. Adjuvant system AS01: helping to overcome the challenges of modern vaccines. Expert Rev Vaccines 16:55–63.

62. Agger EM, Rosenkrands I, Hansen J, Brahimi K, Vandahl BS, Aagaard C, Werninghaus K, Kirschning C, Lang R, Christensen D, Theisen M, Follmann F, Andersen P. 2008. Cationic liposomes formulated with synthetic mycobacterial cordfactor (CAF01): a versatile adjuvant for vaccines with different immunological requirements. PloS one 3:e3116–e3116.

63. Cox MM, Patriarca PA, Treanor J. 2008. FluBlok, a recombinant hemagglutinin influenza vaccine. Influenza Other Respir Viruses 2:211–219.

64. Holtz KM, Robinson PS, Matthews EE, Hashimoto Y, McPherson CE, Khramtsov N, Reifler MJ, Meghrous J, Rhodes DG, Cox MM, Srivastava IK. 2014. Modifications of cysteine residues in the transmembrane and cytoplasmic domains of a recombinant hemagglutinin protein prevent cross-linked multimer formation and potency loss. BMC Biotechnol 14:111.

65. Tian JH, Patel N, Haupt R, Zhou H, Weston S, Hammond H, Logue J, Portnoff AD, Norton J, Guebre-Xabier M, Zhou B, Jacobson K, Maciejewski S, Khatoon R, Wisniewska M, Moffitt W, Kluepfel-Stahl S, Ekechukwu B, Papin J, Boddapati S, Jason Wong C, Piedra PA, Frieman MB, Massare MJ, Fries L, Bengtsson KL, Stertman L, Ellingsworth L, Glenn G, Smith G. 2021. SARS-CoV-2 spike glycoprotein vaccine candidate NVX-CoV2373 immunogenicity in baboons and protection in mice. Nat Commun 12:372.

66. He L, Chaudhary A, Lin X, Sou C, Alkutkar T, Kumar S, Ngo T, Kosviner E, Ozorowski G, Stanfield RL, Ward AB, Wilson IA, Zhu J. 2021. Single-component multilayered self-assembling nanoparticles presenting rationally designed glycoprotein trimers as Ebola virus vaccines. Nat Commun 12:2633.

67. Brouwer PJM, Sanders RW. 2019. Presentation of HIV-1 envelope glycoprotein trimers on diverse nanoparticle platforms. Curr Opin HIV AIDS 14:302–308.

68. Kumar S, Lin X, Ngo T, Shapero B, Sou C, Allen JD, Copps J, Zhang L, Ozorowski G, He L, Crispin M, Ward AB, Wilson IA, Zhu J. 2021. Neutralizing Antibodies Induced by First-Generation gp41-Stabilized HIV-1 Envelope Trimers and Nanoparticles. mBio 12:e0042921.

69. Zhang YN, Paynter J, Antanasijevic A, Allen JD, Eldad M, Lee YZ, Copps J, Newby ML, He L, Chavez D, Frost P, Goodroe A, Dutton J, Lanford R, Chen C, Wilson IA, Crispin M, Ward AB, Zhu J. 2023. Single-component multilayered self-assembling protein nanoparticles presenting glycan-trimmed uncleaved prefusion optimized envelope trimmers as HIV-1 vaccine candidates. Nat Commun 14:1985.

70. He L, Lin X, Wang Y, Abraham C, Sou C, Ngo T, Zhang Y, Wilson IA, Zhu J. 2021. Single-component, self-assembling, protein nanoparticles presenting the receptor binding domain and stabilized spike as SARS-CoV-2 vaccine candidates. Sci Adv 7.

71. Marcandalli J, Fiala B, Ols S, Perotti M, de van der Schueren W, Snijder J, Hodge E, Benhaim M, Ravichandran R, Carter L, Sheffler W, Brunner L, Lawrenz M, Dubois P, Lanzavecchia A, Sallusto F, Lee KK, Veesler D, Correnti CE, Stewart LJ, Baker D, Lore K, Perez L, King NP. 2019. Induction of Potent Neutralizing Antibody Responses by a Designed Protein Nanoparticle Vaccine for Respiratory Syncytial Virus. Cell 176:1420–1431 e1417.

72. Kang HJ, Baker EN. 2011. Intramolecular isopeptide bonds: protein crosslinks built for stress? Trends Biochem Sci 36:229–237.

73. Salas JH, Urbanowicz RA, Guest JD, Frumento N, Figueroa A, Clark KE, Keck Z, Cowton VM, Cole SJ, Patel AH, Fuerst TR, Drummer HE, Major M, Tarr AW, Ball JK, Law M, Pierce BG, Foung SKH, Bailey JR. 2022. An Antigenically Diverse, Representative Panel of Envelope Glycoproteins for Hepatitis C Virus Vaccine Development. Gastroenterology 162:562–574.

74. Chumbe A, Grobben M, Capella-Pujol J, Koekkoek SM, Zon I, Slamanig S, Merat SJ, Beaumont T, Sliepen K, Schinkel J, van Gils MJ. 2024. A panel of hepatitis C virus glycoproteins for the characterization of antibody responses using antibodies with diverse recognition and neutralization patterns. Virus Res 341:199308.

75. Urbanowicz RA, McClure CP, Brown RJ, Tsoleridis T, Persson MA, Krey T, Irving WL, Ball JK, Tarr AW. 2015. A Diverse Panel of Hepatitis C Virus Glycoproteins for Use in Vaccine Research Reveals Extremes of Monoclonal Antibody Neutralization Resistance. J Virol 90:3288–3301.

76. Bailey JR, Barnes E, Cox AL. 2019. Approaches, Progress, and Challenges to Hepatitis C Vaccine Development. Gastroenterology 156:418–430.

77. Fan C, Keeffe JR, Malecek KE, Cohen AA, West AP, Jr., Baharani VA, Rorick AV, Gao H, Gnanapragasam PNP, Rho S, Alvarez J, Segovia LN, Hatziioannou T, Bieniasz PD, Bjorkman PJ. 2025. Cross-reactive sarbecovirus antibodies induced by mosaic RBD nanoparticles. Proc Natl Acad Sci U S A 122:e2501637122.

78. Pierce BG, Boucher EN, Piepenbrink KH, Ejemel M, Rapp CA, Thomas WD, Jr., Sundberg EJ, Weng Z, Wang Y. 2017. Structure-Based Design of Hepatitis C Virus Vaccines That Elicit Neutralizing Antibody Responses to a Conserved Epitope. J Virol 91:e01032–01017.

79. Keck ZY, Wang Y, Lau P, Lund G, Rangarajan S, Fauvelle C, Liao GC, Holtsberg FW, Warfield KL, Aman MJ, Pierce BG, Fuerst TR, Bailey JR, Baumert TF, Mariuzza RA, Kneteman NM, Foung SK. 2016. Affinity maturation of a broadly neutralizing human monoclonal antibody that prevents acute hepatitis C virus infection in mice. Hepatology 64:1922–1933.

80. Broering TJ, Garrity KA, Boatright NK, Sloan SE, Sandor F, Thomas WD, Jr., Szabo G, Finberg RW, Ambrosino DM, Babcock GJ. 2009. Identification and characterization of broadly neutralizing human monoclonal antibodies directed against the E2 envelope glycoprotein of hepatitis C virus. J Virol 83:12473–12482.

81. Keck ZY, Sung VM, Perkins S, Rowe J, Paul S, Liang TJ, Lai MM, Foung SK. 2004. Human monoclonal antibody to hepatitis C virus E1 glycoprotein that blocks virus attachment and viral infectivity. J Virol 78:7257–7263.

82. Grant T, Rohou A, Grigorieff N. 2018. cisTEM, user-friendly software for single-particle image processing. Elife 7.

83. Rames M, Yu Y, Ren G. 2014. Optimized negative staining: a high-throughput protocol for examining small and asymmetric protein structure by electron microscopy. J Vis Exp doi:10.3791/51087:e51087.

84. Punjani A, Rubinstein JL, Fleet DJ, Brubaker MA. 2017. cryoSPARC: algorithms for rapid unsupervised cryo-EM structure determination. Nat Methods 14:290–296.

85. Lebowitz J, Lewis MS, Schuck P. 2002. Modern analytical ultracentrifugation in protein science: a tutorial review. Protein Sci 11:2067–2079.

86. Laue TM, Shah BD, Ridgeway TM, Pelletier SL. 1992. Analytical ultracentrifugation in biochemistry and polymer science. Royal Society of Chemistry.

87. Andrianov AK, Marin A, Wang R, Karauzum H, Chowdhury A, Agnihotri P, Yunus AS, Mariuzza RA, Fuerst TR. 2020. Supramolecular assembly of Toll-like receptor 7/8 agonist into multimeric water-soluble constructs enables superior immune stimulation. ACS Appl Bio Mater 3:3187–3195.

88. Andrianov AK, Marin A, Wang R, Chowdhury A, Agnihotri P, Yunus AS, Pierce BG, Mariuzza RA, Fuerst TR. 2021. In Vivo and In Vitro Potency of Polyphosphazene Immunoadjuvants with Hepatitis C Virus Antigen and the Role of Their Supramolecular Assembly. Mol Pharm 18:726–734.

89. Urbanowicz RA, Wang R, Schiel JE, Keck ZY, Kerzic MC, Lau P, Rangarajan S, Garagusi KJ, Tan L, Guest JD, Ball JK, Pierce BG, Mariuzza RA, Foung SKH, Fuerst TR. 2019. Antigenicity and Immunogenicity of Differentially Glycosylated Hepatitis C Virus E2 Envelope Proteins Expressed in Mammalian and Insect Cells. J Virol 93:e01403–01418.

